# A combined computational fluid dynamics and MRI Arterial Spin Labeling modeling strategy to quantify patient-specific cerebral hemodynamics in cerebrovascular occlusive disease

**DOI:** 10.1101/2021.01.21.426887

**Authors:** Jonas Schollenberger, Nicholas Harold Osborne, Luis Hernandez-Garcia, C. Alberto Figueroa

## Abstract

Cerebral hemodynamics in the presence of cerebrovascular occlusive disease (CVOD) are influenced by the anatomy of the intracranial arteries, the degree of stenosis, the patency of collateral pathways, and the condition of the cerebral microvasculature. Accurate characterization of cerebral hemodynamics is a challenging problem. In this work, we present a strategy to quantify cerebral hemodynamics using computational fluid dynamics (CFD) in combination with arterial spin labeling MRI (ASL). First, we calibrated patient-specific CFD outflow boundary conditions using ASL-derived flow splits in the Circle of Willis. Following, we validated the calibrated CFD model by evaluating the fractional blood supply from the main neck arteries to the vascular territories using Lagrangian particle tracking and comparing the results against vessel-selective ASL (VS-ASL). Finally, cerebral hemodynamics were assessed in two patients with CVOD and a healthy control subject. We demonstrated that the calibrated CFD model accurately reproduced the fractional blood supply to the vascular territories, as obtained from VS-ASL. The two patients revealed significant differences in pressure drop over the stenosis, collateral flow, and resistance of the distal vasculature, despite similar degrees of clinical stenosis severity. Our results demonstrated the advantages of a patient-specific CFD analysis for assessing the hemodynamic impact of stenosis.

## 1. Introduction

Cerebrovascular occlusive disease (CVOD), characterized by the presence of stenosis in the arteries supplying the brain, is a major risk factor for ischemic stroke. Clinical diagnosis and stratification of CVOD patients relies routinely on measuring the maximum narrowing of the lumen based on duplex ultrasound or computed tomography angiography (CTA). However, the degree of luminal stenosis is only one factor in the assessment of stroke risk. Plaque characteristics, downstream brain perfusion, and patency of collateral pathways also play an important role in the overall risk evaluation of cerebral ischemia [1][2][3]. Collateral flow in the circle of Willis (CoW) has been associated with reduced stroke risk in patients with severe carotid stenosis [4][5][6]. Collateral flow is highly dependent on the cerebral vasculature anatomy, availability of collateral pathways, degree of stenosis in the arteries supplying the brain and, critically, the condition of the cerebral microcirculation and its autoregulatory response [7][8].

The clinical gold standard for evaluating collateral flow is digital subtraction angiography (DSA). Despite providing high-resolution images of blood supply in the cerebral arteries, the procedure is invasive and strictly qualtative. MRI arterial spin labeling (ASL) has become an increasingly popular method for measuring cerebral perfusion, and it provides a non-invasive quantitative alternative to DSA. In non-selective ASL (NS-ASL), brain tissue perfusion is measured by magnetically labeling blood in the neck arteries and acquiring a series of slices of the brain after a short transit delay [9]. More recently, ASL has been extended to vessel-selective labeling to measure the perfusion territory of individual arteries [10][11]. The diagnostic capabilities of vessel-selective ASL (VS-ASL) have previously been demonstrated in patients with extracranial stenosis and arteriovenous malformation [12][13]. Additionally, cerebral angiograms have been performed based on VS-ASL to visualize blood supply in the cerebral arteries [14], rendering similar qualitative information on cerebral flow patterns as DSA. Nevertheless, the information provided by VS-ASL on collateral flow patterns has thus far been qualitative.

Image-based computational fluid dynamics (CFD) provides a powerful tool for analyzing cerebral hemodynamics. Compared to experimental approaches, CFD renders velocity, pressure, wall shear stress, etc. throughout entire vascular territories with arbitrarily high spatial and temporal resolutions. The feasibility of CFD to assess cerebral hemodynamics has been previously demonstrated for intracranial stenoses [15][16] and aneurysms [17][18][19]. However, patient-specific calibration of cerebral blood flow CFD models remains challenging. Previous studies have heavily relied on literature data for determining flow splits in the CoW [20][21] or used simplistic allometric scaling assumptions to calibrate outflow boundary conditions [22].

In this paper, we propose a novel strategy to quantitatively characterize regional cerebral blood flow and perfusion using CFD in combination with PC-MRI and ASL data. First, a method to calibrate the cerebral blood flow CFD model based on NS-ASL perfusion data is presented. The calibration includes estimation of flow splits in the CoW from non-selective perfusion images and total inflow to the CoW from PC-MRI, followed by tuning of the outflow boundary conditions to match the estimated flow splits. Second, the calibrated CFD model is validated against territorial perfusion maps from VS-ASL based on the blood supply to each cerebral territory using Lagrangian particle tracking (LPT). Lastly, the proposed strategy is demonstrated via an in-depth quantification of patient-specific cerebral hemodynamics in a healthy control subject and two CVOD patients.

## 2. Material and Methods

### 2.1 Patient details

Two CVOD patients and a healthy control subject were enrolled in a feasibility study and underwent a research MRI exam. The protocol was approved by the local Institutional Review Board and all subjects provided informed written consent (HUM00114275 and HUM00018426). The reconstructed geometric models of the three subjects are illustrated in Fig. 1. The models include the ascending and proximal descending thoracic aorta, its upper branches (brachiocephalic trunk, left carotid and left subclavian), the main neck arteries (internal and external carotids, vertebral arteries), and the main intracranial arteries including the CoW. The healthy control and patient 1 were reconstructed based on magnetic resonance angiography (MRA) and patient 2 based on CTA.

**Fig. 1:**
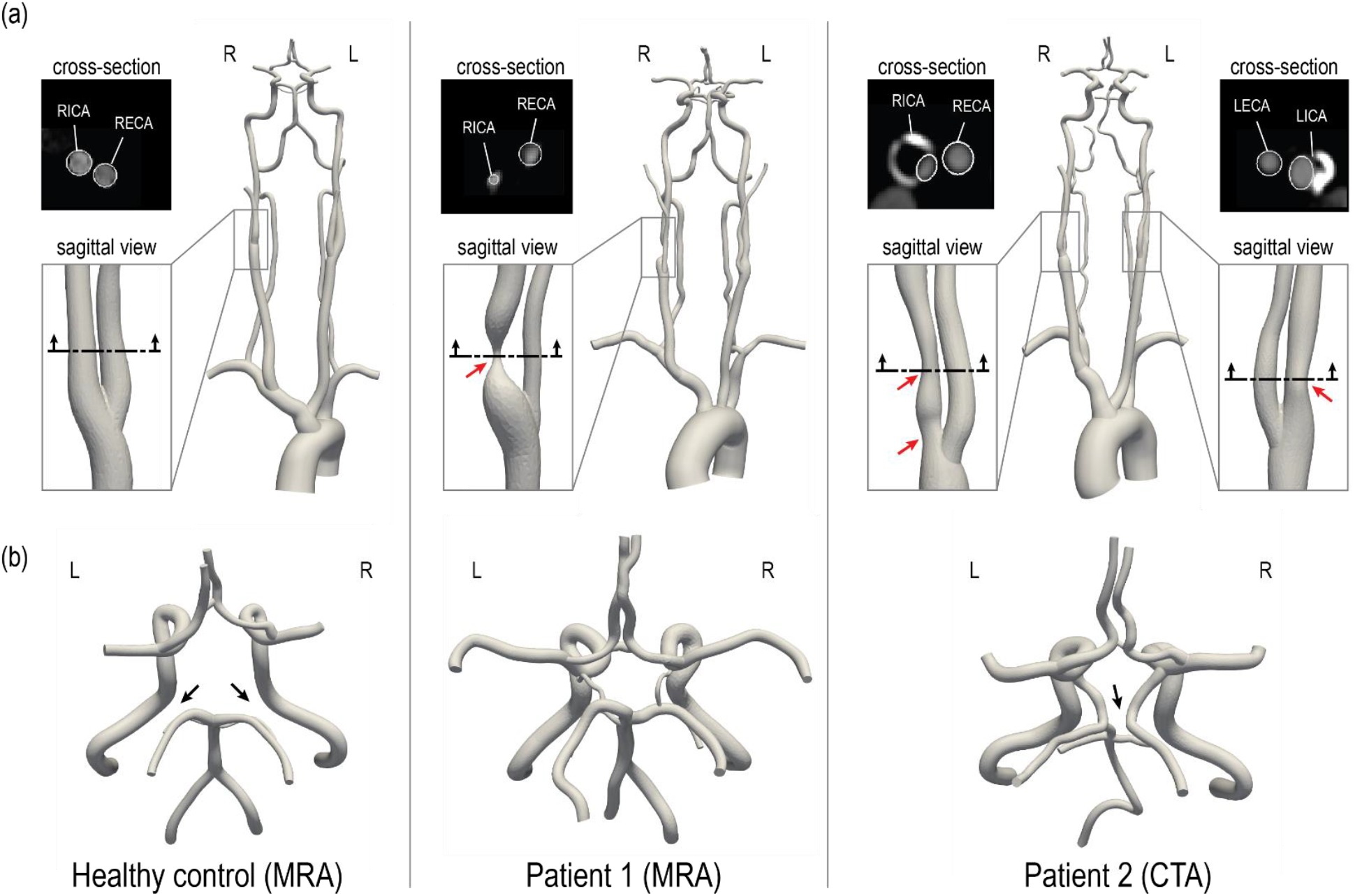
3D-reconstructed geometric models of a healthy control and two CVOD patients. **(a)** For each patient, a close-up of the stenosis is shown. The red arrows indicate the location of the stenosis. An axial cross-section of the stenosis illustrates a comparison between image data and model contours (this comparison is also shown for the healthy volunteer). **(b)** Posterior view of the CoW. The black arrows indicate variations in the CoW anatomy. RICA = right internal carotid artery; RECA = right external carotid artery; LICA = left internal carotid artery; LECA = left external carotid artery. R and L indicate the right and left side from the subject’s perspective.

The healthy control subject (male, 28 y/o) presented without evidence of CVOD. The CoW anatomy was incomplete with both right and left posterior communicating arteries hypoplasia. Patient 1 (female, 55 y/o) presented with an asymptomatic 70-99% stenosis (duplex ultrasound, velocity criteria) in the right proximal internal carotid artery (RICA). The left internal carotid artery (LICA) was patent with no evidence of hemodynamically significant stenosis. Patient 1 has a complete CoW anatomy. Patient 2 (male, 64 y/o) presented with asymptomatic bilateral carotid stenosis. The RICA revealed a tandem stenosis of 80-90% (CTA, ECST criteria), stretching from the carotid bifurcation to the distal end of the carotid bulb. The LICA showed a 60% stenosis (CTA, ECST criteria). Patient 2 has an incomplete CoW anatomy with right P1 segment and distal right vertebral artery (RVA) hypoplasia.

### 2.2 Imaging data

All subjects underwent an MRI study to collect data on vascular anatomy, brain tissue perfusion, and flow. The protocol was performed at 3T (MR750; GE Healthcare, Waukesha, WI) with a 32-channel receive-only head coil for the head and neck and the scanner’s build-in coil for the upper chest. At the end of the study, the subject’s blood pressure was measured in the upper arm in the supine position.

#### Anatomy

Anatomical information from the ascending thoracic aorta to the carotid bifurcation level was acquired with a 2D T1-weighted spoiled gradient echo sequence (voxel size = 0.58 × 0.58 × 2 mm3, TR/TE = 75/3.3 ms). The remaining anatomy from the neck to the CoW was acquired with a 3D Time-of-Flight sequence (voxel size = 0.42 × 0.42 × 1.5 mm^3^, TR/TE = 21/2.5 ms). Additionally, structural images of the brain were collected with a 2D T1-weighted spoiled gradient echo sequence (voxel size = 0.46 × 0.46 × 7 mm3, TR/TE = 100/3.0 ms). For patient 2, an additional CTA data of the neck and head vasculature was available (voxel size = 0.39 × 0.39 × 0.62 mm^3^). For patient 2, the CTA dataset was chosen over the MRA for reconstruction due to the higher resolution.

#### Brain tissue perfusion

Using a pseudo-continuous ASL scheme, non-selective and vessel-selective cerebral perfusion images were collected. Prior to image acquisition, an off-resonance calibration pre-scan was performed to correct for B0-inhomogeneity in the label plane. For the NS-ASL acquisition, sequence parameters were set following consensus recommendations [9]: Label duration = 1800 ms, post-labeling delay = 2000 ms, TR/TE = 4600/4, voxel size = 3.75 × 3.75 × 7 mm^3^, 3D spiral acquisition, 18 slices, 8 pairs of label/control images. The slice prescription was the same as for the T1-weighted structural images. The label plane was positioned above the carotid bifurcation where the arteries of interest (carotid and vertebral) run perpendicular to the plane and with a maximum distance between them. The start of the labeling period was cardiac-triggered to reduce pulsatility artifacts. A proton density image was collected, followed by a non-selective perfusion scan. Subsequently, four VS-ASL scans of the vertebral and carotid arteries were collected. The position of vertebral and carotid arteries within the label plane was determined from the Time-of-Flight acquisition. Keeping all parameters of the NS-ASL scan unchanged, vessel-selective labeling was performed based on a super-selective labeling scheme [10], whereby additional in-plane gradients rotate clockwise every radiofrequency pulse to create a circular labeling spot. Finally, the vessel-selective labeling efficiency was measured by collecting an image 2 cm above the labeling plane 10 ms after labeling for 500 ms. Image reconstruction was performed in MATLAB to a resolution of 128 × 128 using zero-padding in k-space. The ASL perfusion signal was calculated by subtracting label and control images and averaging over all acquired pairs. A detailed explanation of the sequence setup and parameters can be found elsewhere [11].

#### Flow

Volumetric blood flow waveforms were measured using 2D cardiac-gated phase-contrast (PC-MRI) at the level of the ascending aorta (voxel size = 0.58 × 0.58x 5 mm3, TR/TE = 5.2/3.1 ms, velocity encoding = 130 cm/s) and above the carotid bifurcation (voxel size = 0.31 × 0.31x 5 mm3, TR/TE = 6.0/3.7 ms, velocity encoding = 100 cm/s). The slice was positioned perpendicular to the arteries of interest and velocity was encoded in the through-plane direction. PC-MRI data was processed in MATLAB to calculate flow rates.

### 2.3 Computational modeling

The key computational modeling tasks, namely three-dimensional anatomical reconstruction, mesh generation, boundary condition specification, and finite element analysis were performed using the validated open-source computational hemodynamics framework CRIMSON [23].

#### Anatomical reconstruction and mesh generation

3D geometric models of the aorta and head and neck vessels, including the CoW, were reconstructed from the anatomical imaging data. Briefly, centerlines and 2D vessel contours were defined for each vessel of interest. Contours were then lofted to create an analytical representation of each vessel and ultimately define a 3D geometric model of the vasculature [20]. This 3D model was then discretized using linear tetrahedral elements. A mesh-adaptation algorithm [24] was used to refine the mesh locally based on local velocity gradients. The final mesh sizes for healthy, patient 1, and patient 2 consisted of 2.16×10^6^, 1.84×10^6^, and 2.39×10^6^ elements, respectively. Mesh-independence was evaluated for patient 1 by creating an additional highly-refined mesh with 6.86×10^6^ elements, which resulted in a difference of less than 1% for the flow rates at each outlet and a maximum difference of 2 % for peak velocity in the center of the stenosis.

#### Boundary conditions

A pulsatile inflow waveform reconstructed from PC-MRI was mapped to a parabolic velocity profile and prescribed at the inlet of the ascending aorta for each geometric model. Each vessel outlet was coupled to a three-element Windkessel model, which consists of a proximal resistance (*R_p_*), a distal resistance (*R_d_*), and a capacitor (*C*) [25]. For each subject, the total arterial resistance is *R_T_* = *P_mean_*/*Q_T_*, where the mean pressure *P_mean_* = 1/3 *p_systolic_* + 2/3 *p_diastolic_* and *Q_T_* is total cardiac output. The total arterial compliance is *C_T_* = (*Q_T,max_* − *Q_T,min_*)/(*p_systolic_* − *p_diastolic_*) ∗ Δ*t*, where *Q_T,max_* and *Q_T,min_* are maximum and minimum values of aortic inflow, and Δ*t* is the time lapse between these values. Initial estimates for the Windkessel model parameters were obtained by distributing *R_T_* and *C_T_* among the different outlets, to obtain *R_i_* and *C*_i_ for vessel *i* = 1,…,13, as described in [26]. The Windkessel parameters were then iteratively adjusted following the scheme described in detail in section 2.4. Lastly, a no-slip boundary condition was assigned to all vessel walls.

#### Finite Element Analysis

Blood was modeled as an incompressible Newtonian fluid with a dynamic viscosity of 0.004 kg·m^−1^·s^−1^ and a density of 1,060 kg·m^−3^. A stabilized finite-element formulation for the incompressible Navier-Stokes equations was employed to solve for blood flow velocity and pressure in the models [27]. Computations were performed using 80 cores on a high-performance computing cluster. Simulations were run using a time step size of 0.1 ms until cycle-to-cycle periodicity was achieved, typically after 5 cardiac cycles.

### 2.4 Patient-specific calibration of outflow boundary conditions for the CFD models

#### 2.4.1 Calculation of mean flow at model outlets

##### Intracranial arteries

The flow distribution among cerebral and cerebellar arteries was derived from the NS-ASL perfusion images, a cerebral territory atlas, and the total inflow to the brain as illustrated in Fig. 2. First, the NS-ASL perfusion images were mapped into a standardized template space (Montreal Neurological Institute ch2better) using the toolbox SPM12 (Wellcome Trust Center for Neuroimaging, London, UK). Next, the standardized perfusion images were segmented using a vascular territory atlas. The vascular territory atlas was derived from a study by Kim et al. [28], in which the perfusion territories of the main cerebral arteries were mapped based on diffusion-weighted MRI in stroke patients. The atlas was extended to include the cerebellum. This extended vascular territory atlas with a resolution of 370 × 301 × 316 is given as a NIfTI dataset in the supplementary materials. In this work, we assumed the following relationship between eight intracranial arteries and seven vascular territories (see Fig. 2): 1) RACA territory, perfused by the RACA (yellow); 2) LACA territory, perfused by the LACA (magenta); 3) RMCA territory, perfused by the RMCA (green); 4) LMCA territory, perfused by the LMCA (light blue); 5) RPCA territory, perfused by the RPCA (orange); 6) LPCA territory, perfused by the LPCA (dark blue); 7) Cerebellum territory, perfused evenly by RSCA and LSCA (red). A perfusion split psj was then calculated by dividing the sum of perfusion signal in each territory *j* = 1,…,7 by the sum of total perfusion signal over the entire brain. The total mean inflow to the CoW 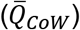 was calculated from the PC-MRI data on left and right ICAs and VAs. Mean flow rates through each intracranial vessel 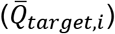 were calculated as the product of the total mean inflow to the CoW and the perfusion split *ps_j_* corresponding to the territory perfused by vessel *i* = 1,…,8.

**Fig. 2:**
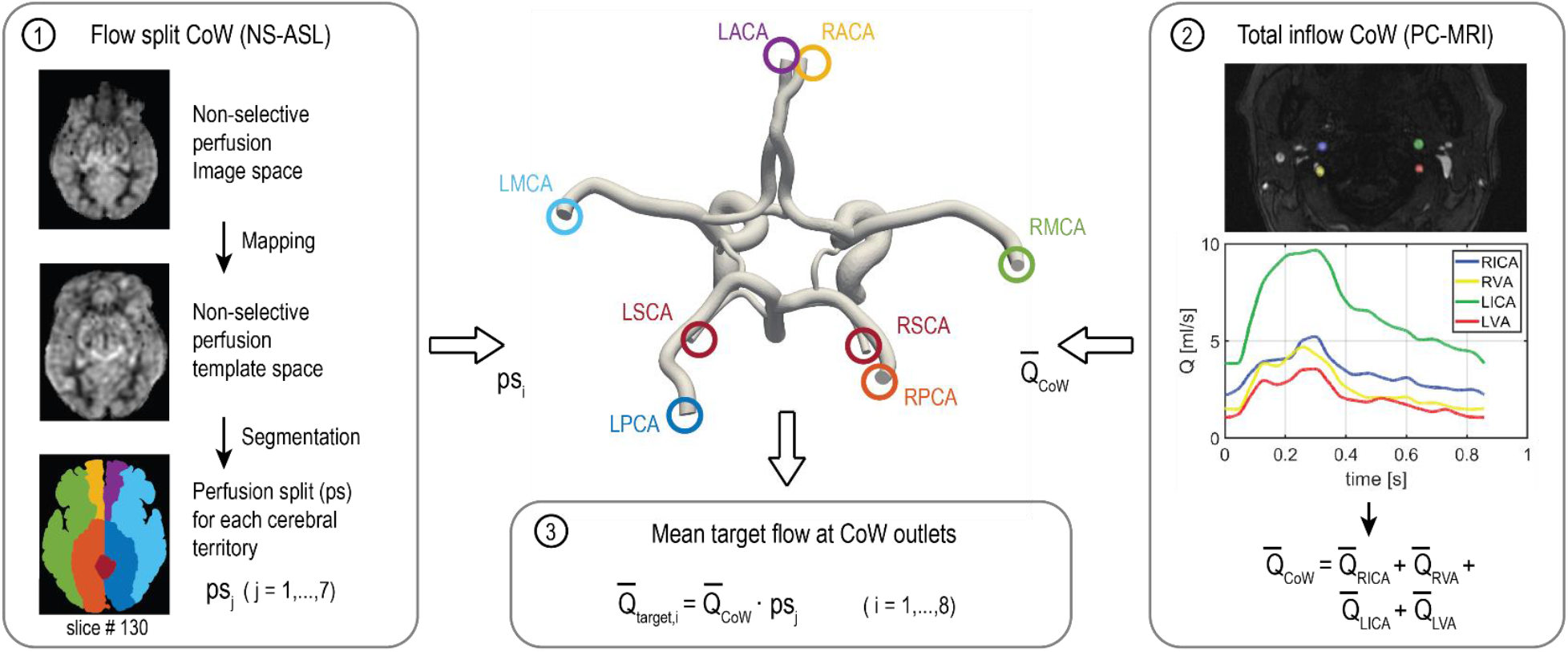
Workflow of calculating the flow split among the arteries of the CoW based on NS-ASL perfusion imaging, a vascular territory atlas, and total inflow to the CoW from PC-MRI. RACA/LACA = right/left anterior cerebral artery, RMCA/LMCA = right/left middle cerebral artery, RPCA/LPCA = right/left posterior cerebral artery, RSCA/LSCA = right/left superior cerebellar artery.

##### Extracranial arteries

Mean flow rates in the external carotid arteries were calculated from PC-MRI. In the subclavian arteries, we assumed a mean flow rate of 5.6% of cardiac-output [20]. Finally, the difference between inflow and intracranial, external carotid arteries, and subclavian arteries flow was assigned to the descending thoracic aorta.

#### 2.4.2 Calibration of Windkessel model parameters

Patient-specific calibration of the Windkessel model parameters for each outflow branch was performed in three stages. *Stage 1*: the distal resistance *R_d_* was iteratively adjusted during simulation runtime using Python controller scripts [29] to match the target mean flow rates. At each simulation time step, *R_d_* was adjusted proportional to the error between the current mean flow and the target flow. Simulations were terminated once the flow at each outlet was fully converged (error < 1%). *Stage 2:* The ratio of *R_p_*/*R_d_* was adjusted for each cerebral and cerebellar branch such that the computed and measured PC-MRI flow waveforms in the ICAs and VAs had similar pulsatility. The total resistance *R_i_* = *R_p_* + *R_d_* at each outlet was kept constant to preserve mean flow. *Stage 3:* Measurements of brachial pressure (*p_systolic_* and *p_diastolic_*) were matched by adjusting *R_T_* and *C_T_*. The percentage change in *R_T_* and *C_T_* through the iterations was proportionally assigned to *R_i_* and *C_i_* at each outlet *i* = 1,…,13. The final Windkessel parameters are summarized in Table S1 of the supplement material.

### 2.5 Validation of the calibrated CFD models

The calibrated CFD models were validated against VS-ASL by comparing the fractional blood supply (FBS) in each vascular territory. We defined the fractional blood supply for a vascular territory *j* from a neck artery *k* as *FBS_j,k_* = *Q_j,k_*/∑_k_ *Q_j,k_*, where *j* = 1,‥,7 is the vascular territory index and *k* = 1,‥,4 is the neck artery index (1:RIVA, 2:RVA, 3:LVA, 4:LICA), and *Q_j,k_* is the flow contribution from the neck artery *k* to the vascular territory *j*. The process for calculating *FBS_j,k_* from VS-ASL and CFD data is described next.

#### 2.5.1 Fractional blood supply based on VS-ASL

The process for calculating *FBS_j,k_* from VS-ASL images is outlined in Fig. 3. The perfusion signal in the VS-ASL images is determined by the blood supply from a single neck artery to the vascular territories. The signal in the VS-ASL images was first scaled based on the measured labeling efficiency. Then, scaled VS-ASL images were transformed into the standardized template space (panel 1). The sum of all scaled VS-ASL images produces the total perfusion image. The *FBS_j,k_* maps were then calculated by dividing each scaled VS-ASL image by the total perfusion image on a voxel-by-voxel basis (panel 2). Then, the *FBS_j,k_* maps were segmented into different territories using the vascular territory atlas. Lastly, due to noise in the raw ASL data, negative values of *FBS_j,k_* distribution are possible on a given voxel. Therefore, we characterize the *FBS_j,k_* distribution by its median (M) and median absolute deviation (MAD) (see panel 3).

**Fig. 3:**
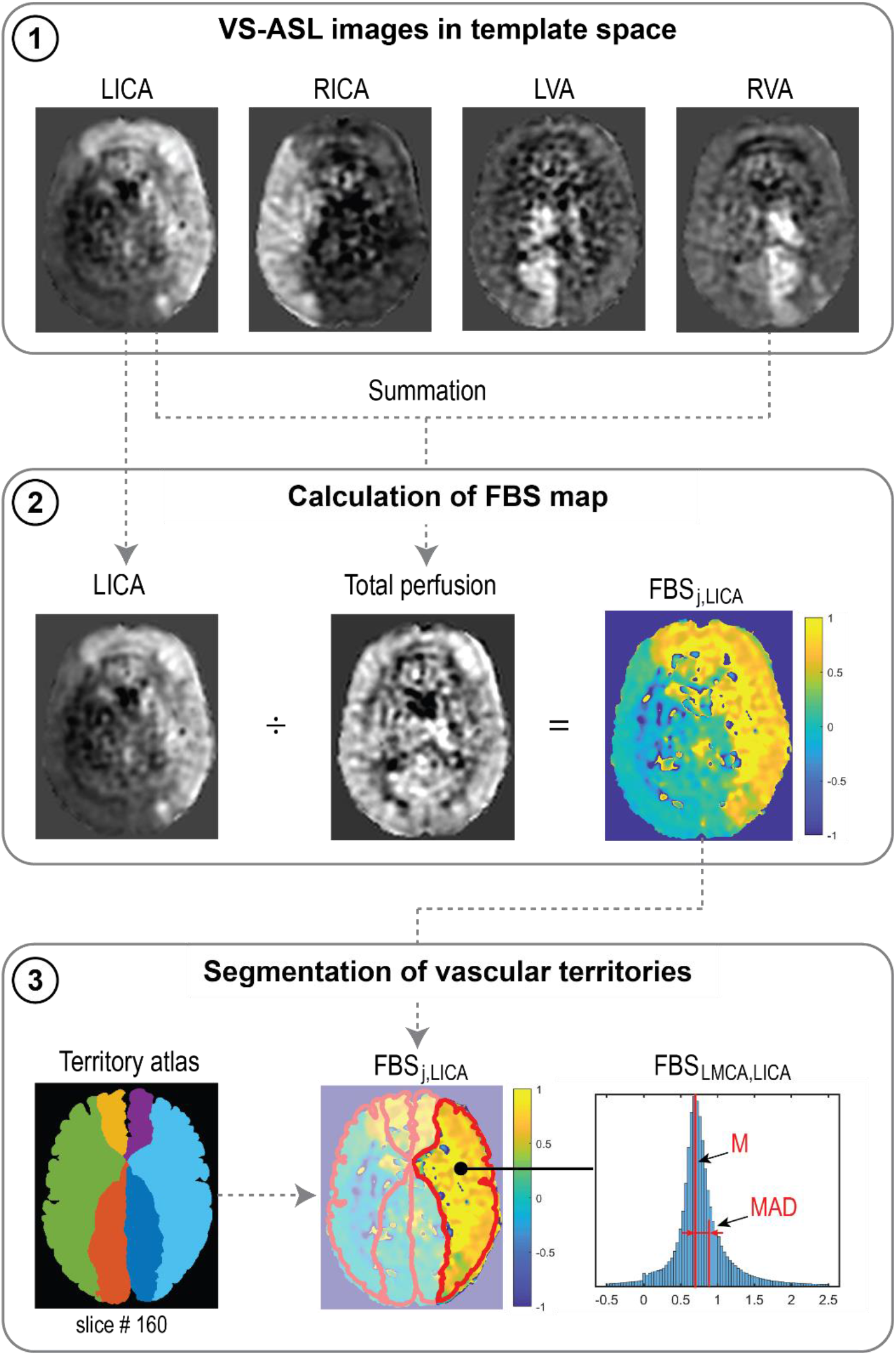
Processes to calculate *FBS_j,k_* from VS-ASL images, illustrated for the LICA. (1) Scaled VS-ASL perfusion images in standardized template space. (2) Calculation of spatial *FBS_j,k_* maps. (3) Segmentation of territories using a vascular territory atlas and calculation of median (M) and median absolute deviation (MAD) from the *FBS_j,k_* distribution in each territory.

#### 2.5.2 Fractional blood supply based on CFD Lagrangian particle tracking

To validate the CFD results using the VS-ASL data, one must develop a method to assess the fractional blood supply of the intracranial arteries/territories and compare it with the median (M) of the *FBS_j,k_* obtained with VS-ASL for each artery/territory. This can be achieved via a further post-processing analysis on the CFD data, known as “Lagrangian particle tracking” (LPT), whereby virtual boluses of blood are created by seeding mass-less particles in different regions of the vasculature and advected by the velocity field over multiple cardiac cycles. This is an established technique with multiple applications in cardiovascular flows [30] [31] [32] [33] [34].

In this work, particles were seeded continuously at the base of the carotid and vertebral arteries (supplement material Fig. S1). LPT was performed individually for each carotid and vertebral artery over 4 cardiac cycles. The number of particles seeded in each artery, over multiple re-injections over the 4 cardiac cycles, was proportional to the flow rate of each artery. Particles were then counted at each outlet of the intracranial arteries, and once cycle-to-cycle periodicity in the number of tracked particles was achieved, the total number of particles collected per vessel over a full cardiac cycle was extracted. Lastly, for each intracranial vessel, assigned to a vascular territory *j*, its fractional blood supply *FBS_j,k_* from each neck artery *k* was calculated by dividing the particle count of the LPT analysis for neck artery *k* by the sum of the particle counts of each of the four LPT analyses of the neck arteries.

## 3. Results

### 3.1 Validation of calibrated CFD model

#### 3.1.1 Fractional blood supply: CFD LPT versus VS-ASL

##### Qualitative analysis

A qualitative comparison between FBS obtained from VS-ASL and CFD LPT is illustrated in Fig. 4. The VS-ASL images (Fig. 4a) show the perfusion territories of the four neck arteries, from the inferior region of the cranium (bottom row slices) to the superior region (top row slices). Each image voxel was color-coded based on the FBS of the neck arteries. For visualization purposes, we limited the FBS in each voxel to positive fraction values between 0 and 1 (cf. Fig. 3 panel 3). LPT analyses results (Fig. 4b) show maps of the advection of particles, color-coded based on the seeding artery in the neck, as well as temporal histograms of particles collected at the outflow of selected intracranial arteries.

1. Middle cerebral arteries: The VS-ASL data revealed that the perfusion territories of the RMCA (arrow 1) and LMCA (arrow 2) were primarily supplied by the ipsilateral carotid artery for all subjects regardless of the degree of stenosis. This perfusion pattern was also replicated in the LPT analyses, where particles exiting the RMCA primarily originated in the RICA (green particles) and particles leaving the LMCA predominantly originated in the LICA (blue particles).
2. Anterior cerebral arteries: All three subjects displayed flow compensation from the LICA to the RACA territory (arrow 3) in the VS-ASL images. The LPT analysis also reproduced this flow compensation in the RACA via the anterior communicating artery (AComA) for all subjects. The histograms of LPT RACA particles illustrate clear differences in the amount of compensation among subjects. The healthy control subject revealed significant blood supply from the LICA (blue particles) to the RACA, despite the absence of stenosis. The severe RICA stenosis in patient 1 resulted in the RACA being predominantly supplied by the LICA. In contrast, the severe stenosis in the RICA and the mild stenosis in the LICA in patient 2 only led to a small amount of collateral flow in the RACA.
3. Posterior cerebral arteries: The VS-ASL images revealed large differences in blood supply in the RPCA (arrow 4) and LPCA (arrow 5) territories between subjects. These differences in blood supply were also replicated in the LPT analyses. In the healthy control subject, the posterior circulation received mixed supply from both vertebral arteries as seen in the LPCA histogram. In patient 1, the RPCA territory was predominantly perfused by the LVA whereas the LPCA territory was predominantly perfused by the RVA (see histogram). The VS-ASL data revealed a switch in blood supply to the posterior circulation between right and left hemisphere, a switch also mirrored in the LPT analysis which shows vortex-like flow patterns in the basilar artery resulting in crossing of particles originating in the VAs. In patient 2, the posterior circulation was supplied by the ipsilateral carotid arteries. In this patient, the VAs did not contribute to cerebral blood flow. Instead, the LVA supplied most of the cerebellum flow with some small contribution from the LICA, as also apparent in the VS-ASL data (arrow 6).

**Fig. 4:**
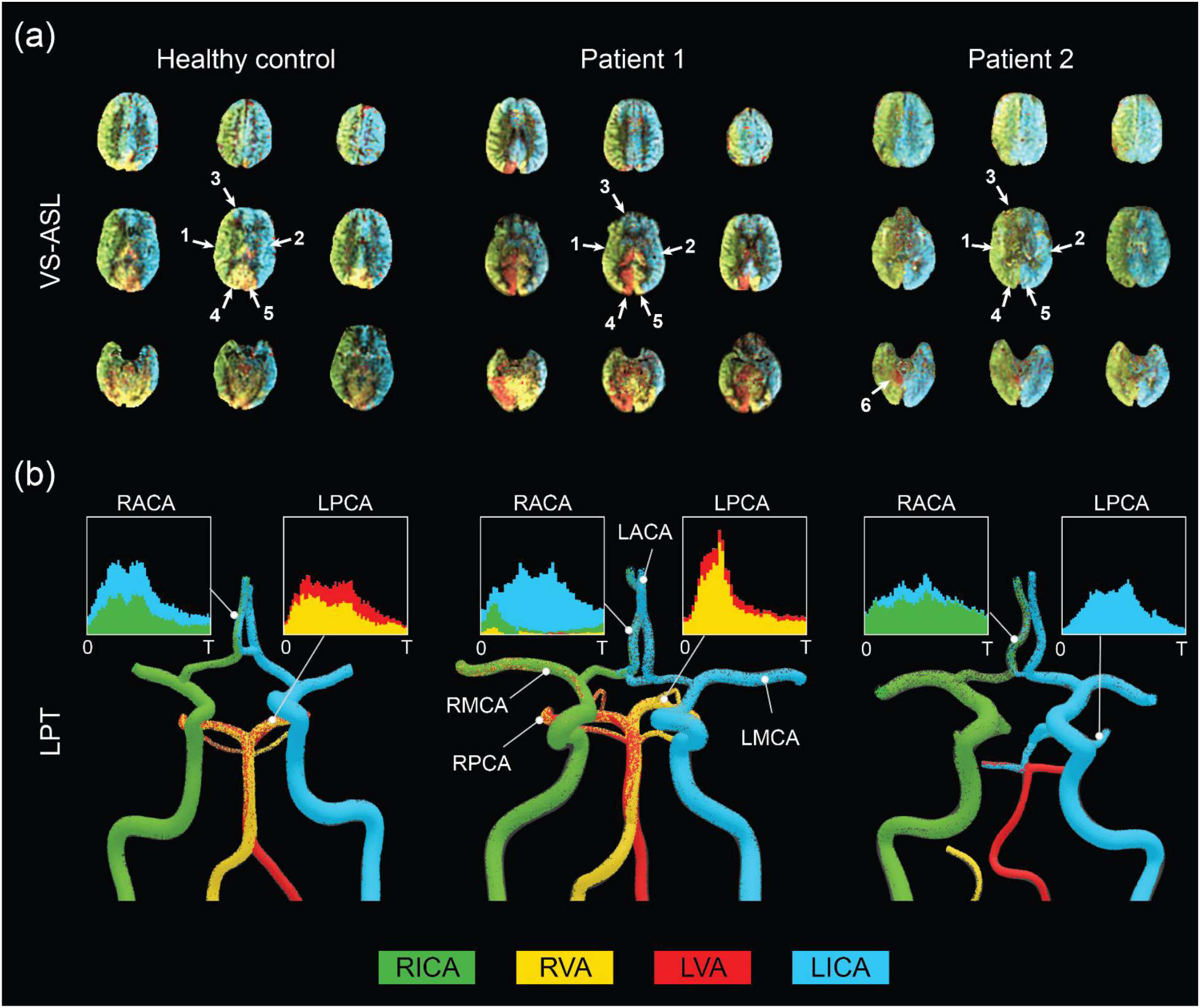
Qualitative comparison between FBS obtained from VS-ASL and CFD LPT **(a)** VS-ASL images show the perfusion territories of the main neck arteries from the inferior of the cranium (bottom row slices) to the superior (top row slices). The images were created by color-coding the FBS maps of the main neck arteries on a voxel-by-voxel basis. For visualization purposes, we limited the fractional contributions of each neck artery to a positive range between 0 and 1. The arrows indicate the vascular territories of the 1) RMCA, 2) LMCA, 3) RACA, 4) RPCA, 5) LPCA, and 6) cerebellum. **(b)** LPT analyses show the advection of particles in the large arteries of the CoW. Particles are color-coded based on the artery of origin in the neck. Histograms demonstrate mixed supply in the RACA and LPCA over the cardiac cycle T.

##### Quantitative analysis

A quantitative comparison of *FBS_j,k_* obtained with VS-ASL and CFD LPT is summarized in Fig. 5. For each vascular territory *j*, the percentage supply contributions from the neck arteries *k*, obtained from VS-ASL (red bars) and LPT (blue bars), are shown. VS-ASL data includes the median absolute deviation. Due to noise in the VS-ASL signal, FBS results show small negative values in territories for which the perfusion contribution from a given neck artery *k* is small. Conversely, the LPT data is “noise-free” and given by a single value of FBS, instead of by a distribution. Overall, VS-ASL and LPT estimates of *FBS_j,k_* agreed well for all subjects. The LPT analysis correctly identified the artery contributing the largest % of perfusion in vascular territories predominately perfused by a single neck artery in all subjects (e.g. LACA, LMCA, RMCA). Furthermore, the magnitude of flow compensation from the LICA to the RACA territory was correctly reflected in the LPT analyses for all subjects. The main sources of perfusion in the RPCA and LPCA territories were correctly identified by the LPT in both patients and only partially matched in the healthy control subject.

**Fig. 5:**
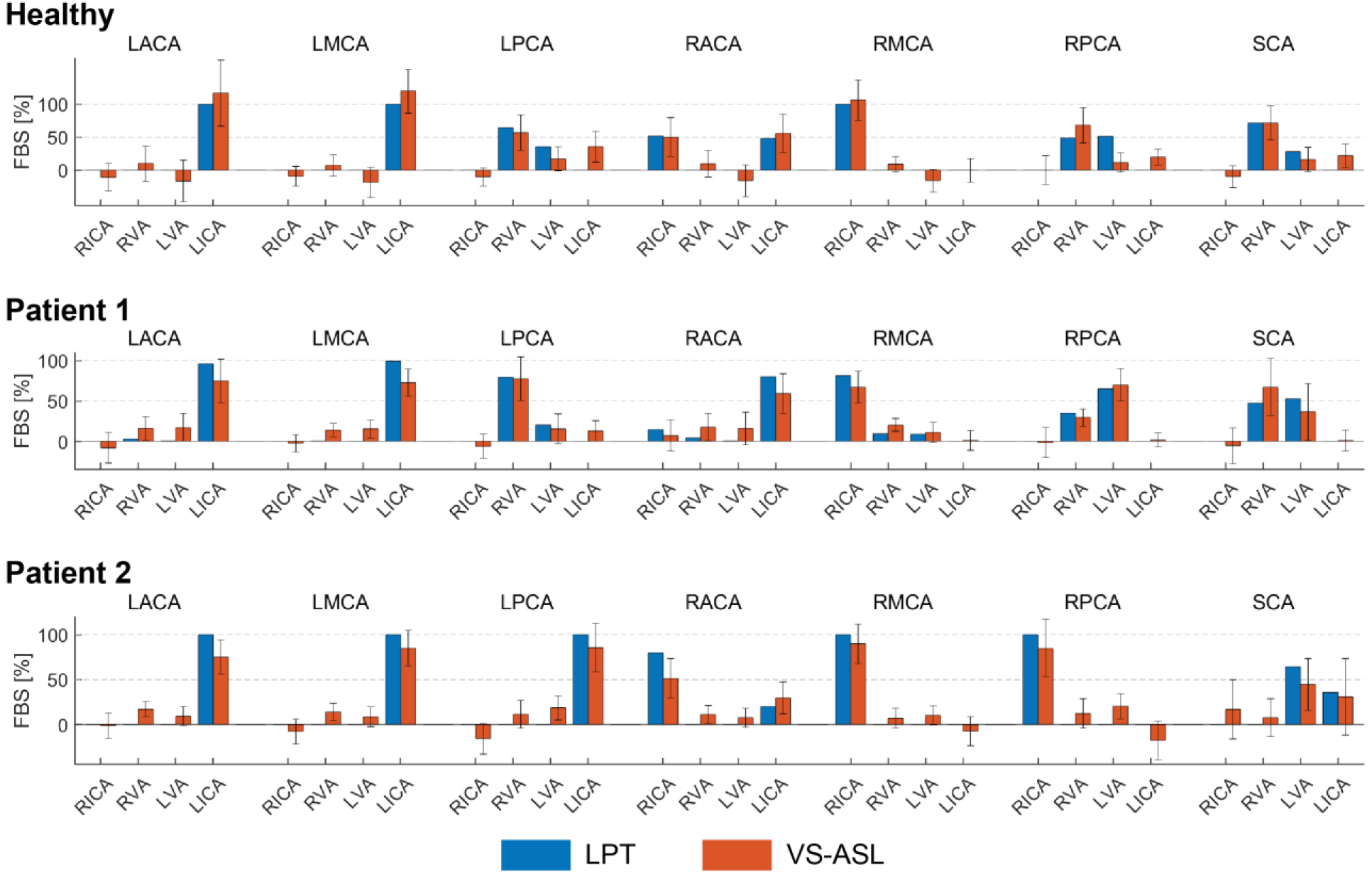
Comparison of the *FBS_j,k_*, obtained from VS-ASL and LPT, in each vascular territory *j* and for each neck artery *k*. For VS-ASL, values of *FBS_j,k_* represent the median of the *FBS_j,k_* distribution in each vascular territory. The error bar represents the median absolute deviation. For LPT, values of *FBS_j,k_* were calculated based on the particle count at each outlet of the CoW. RACA/LACA = right/left anterior cerebral artery, RMCA/LMCA = right/left middle cerebral artery, RPCA/LPCA = right/left posterior cerebral artery, SCA = superior cerebellar arteries, RVA/LVA = right/left vertebral artery, RICA/LICA = right/left internal carotid artery.

A correlation coefficient of *FBS_j,k_* between VS-ASL and LPT was calculated for each subject over all vascular territories *j* and neck arteries *k* (supplement material Fig. S2). The correlation coefficients were *R* = 0.92, *R* = 0.94, and *R* = 0.95 for the healthy subject, patient 1, and patient 2, respectively.

#### 3.1.2 Flow: CFD versus PC-MRI

Mean flow rates from the calibrated CFD model were compared to PC-MRI flow data in the vertebral and carotid arteries above the carotid bifurcation. The difference in mean flow rates in each neck artery was smaller than 10% for all subjects. A comparison of the flow waveforms in the vertebral and carotid arteries between CFD and PC-MRI is shown in Fig. 6 for patient 1, Fig. 7 for patient 2, and Fig. S3 for the healthy control subject.

**Fig. 6:**
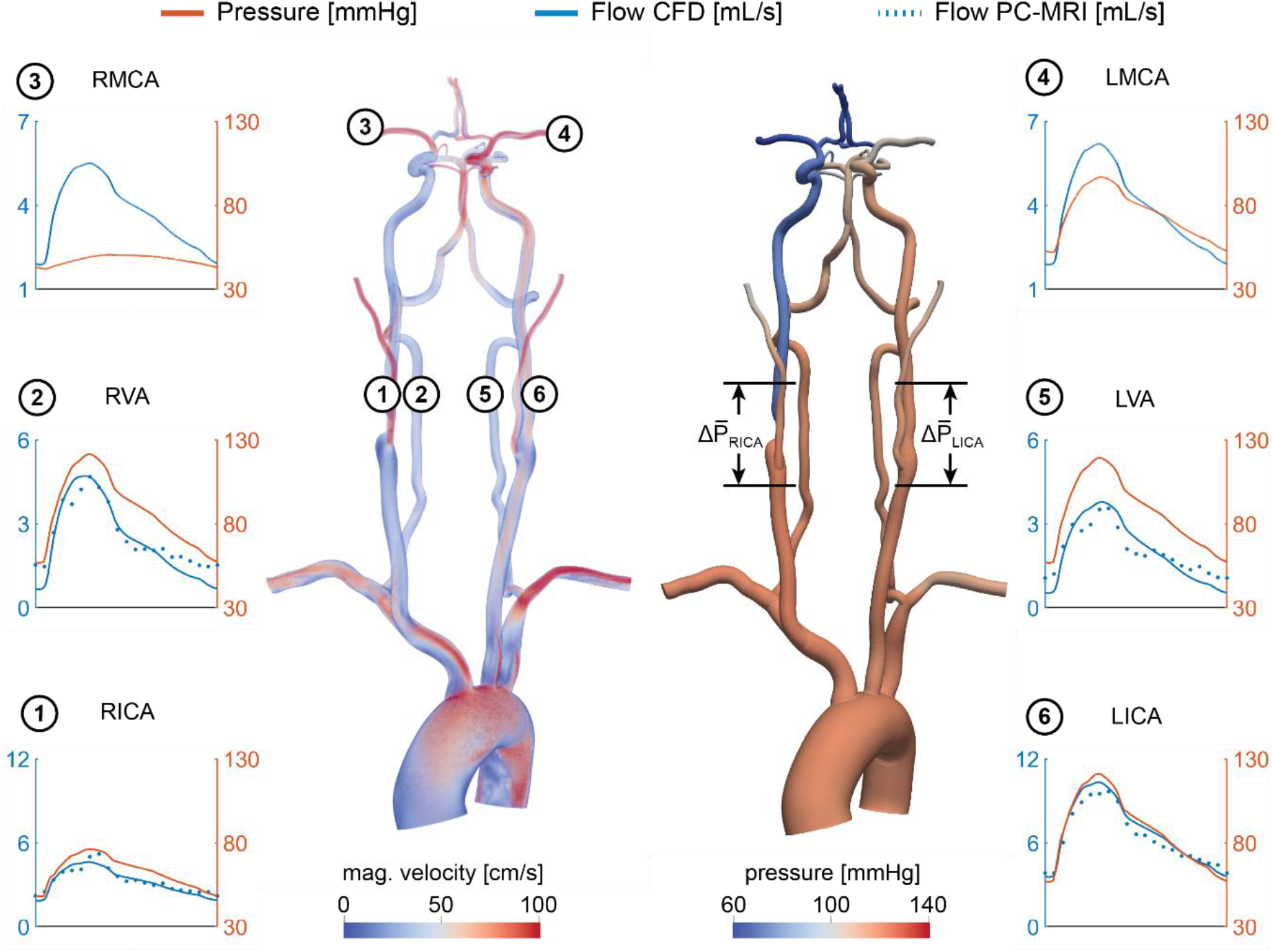
Velocity and pressure fields at peak systole for patient 1. Flow and pressure waveforms are evaluated in the internal carotid (ICA), vertebral (VA), and middle cerebral arteries (MCA). The flow waveforms in the neck arteries are compared to PC-MRI measurements above the carotid bifurcation (dotted lines). The mean pressure drop 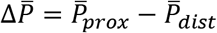 was calculated over the RICA stenosis and the same vessel segment in the LICA.

**Fig. 7:**
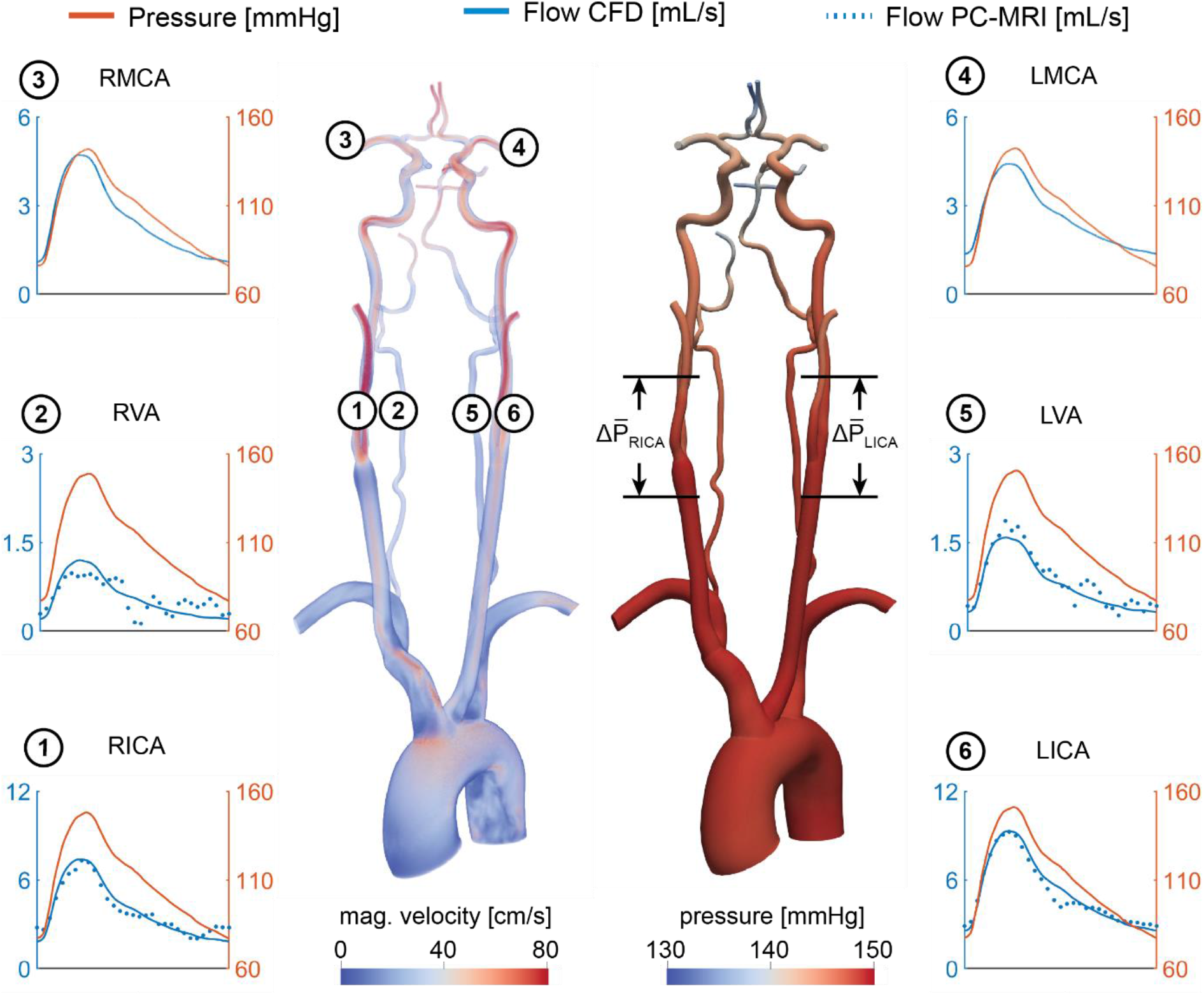
Velocity and pressure fields at peak systole for patient 2. Flow and pressure waveforms are evaluated in the internal carotid (ICA), vertebral (VA), and middle cerebral arteries (MCA). The flow waveforms in the neck arteries are compared to PC-MRI measurements above the carotid bifurcation (dotted lines). The mean pressure drop 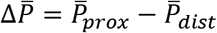 was calculated over the RICA and LICA stenosis.

### 3.2 CFD-based quantification of patient-specific cerebral hemodynamics

Once the validation of the CFD results using VS-ASL and PC-MRI data is established, we use the calibrated models to assess alterations in cerebral hemodynamics. Specifically, pressure and flow waveforms, the hemodynamic impact of the carotid stenoses, and the resistances of the distal cerebral vascular territories were quantified in the two patients of the study.

Fig. 6 shows pressure and flow waveforms in six arteries of the neck and head for patient 1. The mean pressure drop over the stenosis was defined as 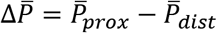, where 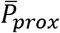 and 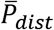 are the mean cross-sectional-averaged pressures 2 cm proximal and distal to the maximum diameter reduction, respectively. The mean pressure drop over the RICA was 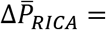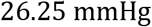, compared to a 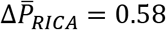 mean pressure drop over the unstenosed LICA.

Another metric of the hemodynamic significance of the stenosis is given by the fractional flow (FF) index [3][15][35], defined as 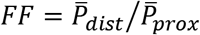. This index produced *FF_RICA_* = 0.71 and *FF_LiCA_* = 0.99. The stenosis resulted in a substantial difference in flow between the RICA and LICA, with mean flow rates of 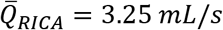 and 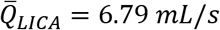. In the CoW, the RMCA and LMCA exhibit substantially different mean values of pressure 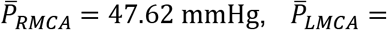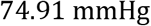, mean difference: 27.29 mmHg). Despite the pressure difference between hemispheres, the mean flow rates in the RMCA and LMCA were comparable with 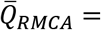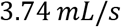 and 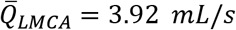. This preservation of flow at the RMCA was achieved through a reduction in the total resistance of its distal vasculature, resulting in *R*_*RMCA*_ = 1.7 ∙ 10^9^ *pa s m*^−3^ compared to *R_LMCA_* = 2.54 ∙ 10^9^ *pa s m*^−3^ at the contralateral LMCA.

Fig. 7 shows the pressure and flow waveforms for patient 2. The mean pressure drop over the stenoses and *FF* indices in the RICA and LICA were 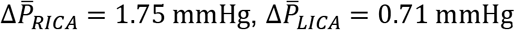, *FF_RICA_* = 0.98, and *FF_LiCA_* = 0.99, respectively. The mean flow rates in the RICA and LICA were 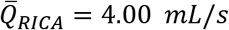 and 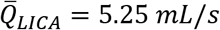. Despite the severe RICA stenosis and moderate LICA stenosis, the mean pressure at the RMCA and LMCA were similar (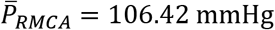,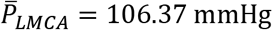, mean difference: 0.05 mmHg). The mean flow rates, as well as the corresponding total resistances, at the outlets of the RMCA and LMCA were comparable with 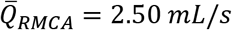 and 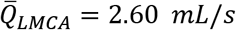, and *R*_*RMCA*_ = 5.64 ∙ 10^9^ *pa s m*^−3^ and *R_LMCA_* = 5.44 ∙ 10^9^ *pa s m*^−3^, respectively.

Pressure and flow waveforms for the healthy control subject are provided in the supplement material (Fig. S3).

## 4. Discussion

### 4.1 Patient-specific calibration of outflow boundary conditions

We presented a strategy for calibrating patient-specific outflow boundary conditions in the CoW. Our strategy relies on deriving mean flow splits in the main arteries of the CoW using ASL perfusion data, and on knowledge of the total flow to the head given by PC-MRI data in the neck arteries. The perfusion data in the different cerebral territories is the result of the spatial distribution of blood supply to the brain tissue, which is determined by the overall distal microvascular resistance and potentially cerebral auto-regulatory effects that seek to compensate deficits in flow in a region of the brain. Therefore, the calibrated outflow boundary condition parameters include the effect of all these mechanisms. Cerebral auto-regulatory compensation was apparent in patient 1, where the ASL-derived flow rates in the right and left MCA of the CoW were comparable despite a severe RICA stenosis. To match the ASL-derived mean flow rates in the CFD analysis, the distal resistance of each Windkessel model for the vessels in the CoW was iteratively adjusted during stage 1 of the calibration (cf. section 2.4.2). The calibrated RMCA resistance was significantly lower than its LMCA counterpart. This finding points to a substantial vasodilation of the distal vasculature of the RMCA to maintain adequate blood supply to the brain tissue.

Previous CFD modeling studies of cerebral blood flow have relied primarily on assumptions on the flow distribution in the CoW, either based on literature data of healthy vasculatures [20][21] or allometric scaling laws [22]. However, in situations of cerebrovascular disease, the distribution of flow between the different vessels of the CoW may be substantially different from that given by idealized allometric scaling principles based on healthy data. Ultimately, incorrect values of flow in the vessels of the CoW will affect the quality of the CFD results. Zhang et al. [36] previously presented a calibrated 1D-0D computational model of cerebral blood using single photon emission computed tomography (SPECT) to estimate the flow distribution in the CoW [37]. In this work, we built on this approach by acquiring non-invasive and non-radioactive NS-ASL perfusion images and using 3D models of blood flow, given by the incompressible Navier-Stokes equations, which are essential to capture complex hemodynamics around the stenosis and in the small and tortuous vessels of the CoW.

### 4.2 Validation of calibrated CFD model

In this work, we demonstrated that our CFD calibration strategy of using NS-ASL perfusion images, in combination with a vascular territory atlas and PC-MRI, can accurately characterize flow in the main arteries of the CoW in patients with CVOD. The LPT analysis performed using the calibrated CFD model was validated using VS-ASL data by comparing their respective values of FBS in each vascular territory. Results showed an overall good agreement between LPT and VS-ASL with high correlation coefficients.

Bockman et al. [22] developed a CFD model of the vertebrobasilar system with outflow boundary conditions defined via allometric scaling on healthy subjects without flow-altering CVOD. They studied the agreement in the laterality of the VA blood supply for the cerebral and cerebellar circulations between CFD and vessel-encoded ASL. ASL perfusion data was therefore not used to calibrate the outflow BC of the CFD model. Here, we modeled the entire CoW and validated blood supply to each vascular territory in both healthy and CVOD subjects. Furthermore, we demonstrated that the calibrated CFD models captured the collateral flow observed in the VS-ASL data.

### 4.3 Assessment of patient-specific cerebral hemodynamics

Using the validated CFD model, we performed an in-depth quantification of cerebral hemodynamics in the two CVOD patients. Both patients presented with a severe RICA stenosis (70-99% diameter reduction) according to the velocity criteria (Patient 1), which correlates the peak systolic velocity measured with Duplex Ultrasound to a percentage diameter reduction, and the ECST criteria (Patient 2), which is defined as the diameter reduction relative to the original vessel diameter based on CTA. Despite similar degrees of clinical stenosis severity, the cerebral hemodynamics varied significantly between these two patients. In patient 1, the RICA stenosis led to severe ipsilateral pressure and flow drop compared to the contralateral unstenosed LICA. Despite the pressure difference between right and left hemisphere, there was no significant difference in flow between the right and left MCA due to collateral flow compensation and vasodilation of the distal vasculature. In contrast, the RICA stenosis in patient 2 did not result in a notable drop in ipsilateral pressure or flow. Consequently, the flow compensation between hemispheres was small. The difference in flow compensation between the two patients is explained by the much smaller stenosis diameter of patient 1 (1.4 mm in our geometric model, 74.7% diameter reduction relative to the distal diameter) compared to patient 2 (3.0 mm, 52.3% diameter reduction relative to distal diameter), see Fig. 1. Flow compensation is highly dependent on accurate characterizations of the degree of stenosis, the cerebral anatomy, and the cerebrovascular reserve and can vary significantly between patients.

Medical imaging used for cerebral hemodynamics assessment (e.g. Transcranial Doppler, 4D Flow MRI, etc.) only provide information on velocity. However, the above results (similar cerebral flow between the two subjects while having substantial differences in pressure) highlight the shortcomings of describing the hemodynamic significance of CVOD lesions purely from the perspective of velocity. In contrast, blood pressure is highly sensitive to changes in vascular resistance induced by the stenosis, as illustrated in patient 1. Therefore, substantial changes in cerebral blood pressure may be a more sensitive marker for diminished vascular flow reserve. While pressure catheter measurements are commonly acquired in other vascular territories (e.g. coronary arteries), cerebral blood pressure is generally not acquired during clinical assessment of CVOD patients due to increased stroke risk. While there have been efforts to derive pressure gradients from 4D Flow MRI data, application in the small and tortuous CoW arteries remains challenging due to limited spatial resolution [38]. In contrast, patient-specific CFD overcomes these shortcomings by providing highly resolved velocity and pressure.

To quantify the stenosis hemodynamic significance, we calculated the fractional flow index, defined as the ratio of the pressures distal and proximal to the stenosis under baseline flow conditions (e.g., non-hyperemic). The fractional flow in the RICA stenosis was *FF_RICA_* = 0.71 for patient 1 and *FF_RICA_* = 0.98 for patient 2. Using a threshold of *FF* = 0.8 [35], only the RICA stenosis in patient 1 would be deemed to be hemodynamically significant. While the clinical metric of diameter reduction resulted in a similar value of 70-99% in the RICA for both patients, the fractional flow index captured better the large differences in cerebral hemodynamics between the patients. The metric of fractional flow reserve (FFR) has become widely used in coronary artery disease (CAD) to evaluate the risk for myocardial ischemia. FFR-guided intervention has been shown to reduce myocardial ischemia, rate of death, and revascularization compared to anatomical-based intervention [39][40]. However, the use of fractional flow for the risk assessment of ischemic stroke in CVOD has not yet been established.

FBS obtained from CFD and LPT provided important information about collateral flow in the anterior circulation and mixed VA supply in the posterior circulation. Beyond flow compensation, FBS in the cerebral vascular territories can provide clinically relevant information about the etiology of embolic stroke. Even after thorough diagnostic evaluation, the cause of embolic stroke remains uncertain in one third of cases [41]. It is common that multiple atherosclerotic lesions within the same patient are identified. Evaluation of FBS in the region of the stroke could help to determine the source of emboli and ultimately guide clinical treatment.

These results illustrate the level of insight on hemodynamic assessment of CVOD patients that calibrated patient-specific CFD analysis can bring.

### 4.4 Limitations

#### Vascular territory atlas

Estimating the flow splits between the main arteries of the CoW and, consequently, calibrating the corresponding outflow Windkessel models, relies on the segmentation of non-selective ASL perfusion images using a vascular territory atlas. The atlas was derived from a large population study and represents a map of the average vascular territories in the brain. Although the size of border zones between territories appears to be much narrower than previously assumed [28], a degree of variability in the vascular territories between patients is expected. Furthermore, leptomeningeal collaterals can form between the distal branches of the CoW to augment flow compensation in cases of severe stenosis [42], thereby altering the vascular territories of the main cerebral arteries. To assess the accuracy of the atlas in the subjects of our study, we could qualitatively compare their boundaries against the outlines of the perfusion VS-ASL data, see Fig. 3, maps of territorial atlas and *FBS_j,LiCA_* for slice 160.

#### VS-ASL

Perfusion quantification from ASL images is challenged by its inherently low signal-to-noise ratio, pulsatility artifacts, and sensitivity to bolus arrival time variation. Although NS-ASL is rapidly gaining acceptance in the clinic, quantification of perfusion is not readily available in all scanner platforms and the technique is generally used qualitatively in the clinical setting [9]. In VS-ASL, these challenges can lead to artifacts in the territorial perfusion maps [11]. For example, signal fluctuations between label and control images can result in spurious perfusion signal in vascular territories not perfused by the labeled neck artery, and consequently lead to errors in the calculation of the FBS. To account for the spatial variability of FBS within a vascular territory, we included the median absolute deviation in the data as shown in Fig. 5.

## 5. Conclusion

In this work, we presented a strategy to quantify cerebral hemodynamics using CFD in combination with ASL and PC-MRI data. We demonstrated that our calibrated CFD model accurately reproduced the fractional blood supply to the vascular territories, as obtained from VS-ASL. In particular, the flow compensation between hemispheres was captured well by the calibrated CFD models. The assessment of cerebral hemodynamics in two CVOD patients using calibrated patient-specific CFD analysis showed significant differences in cerebral hemodynamics between the patients despite similar degrees of clinical stenosis severity. We further illustrated the advantages of CFD-based pressure data for assessing the hemodynamic significance of carotid stenosis. Future studies are needed to investigate the benefits of using of a hemodynamic-based metric (fractional flow) versus an anatomy-based metric (diameter reduction) for the risk assessment of CVOD patients.

## 6. Acknowledgements

The authors would like to acknowledge Heather Golden for her help with patient enrollment.

## 7. Author’s contributions

<u>Jonas Schollenberger</u>: Conceptualization of the study, patient enrollment, data acquisition, data analysis and interpretation, development of calibration strategy, development of post-processing routines, computational simulation, preparation of manuscript

<u>Nicholas H. Osborne</u>: patient enrollment, clinical supervision, data interpretation, review of manuscript

<u>Luis Hernandez-Garcia</u>: Conceptualization of the study, data acquisition, review of manuscript, supervision

<u>C. Alberto Figueroa</u>: Conceptualization of the study, data interpretation, preparation of manuscript, funding, supervision

## 10. Supplement material

Supplemental material for this article is available online.

## 13. Supplement material

**Fig. S1:**
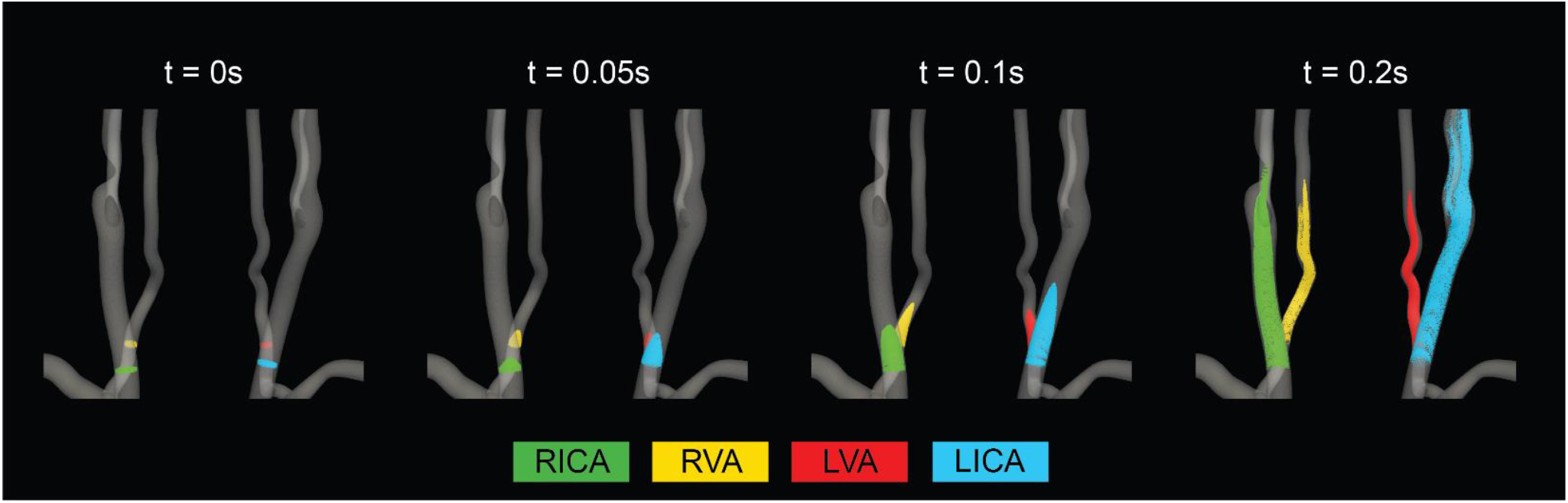
Continuous seeding and advection of Lagrangian particles in the vertebral and carotid arteries at four time points.

**Fig. S2:**
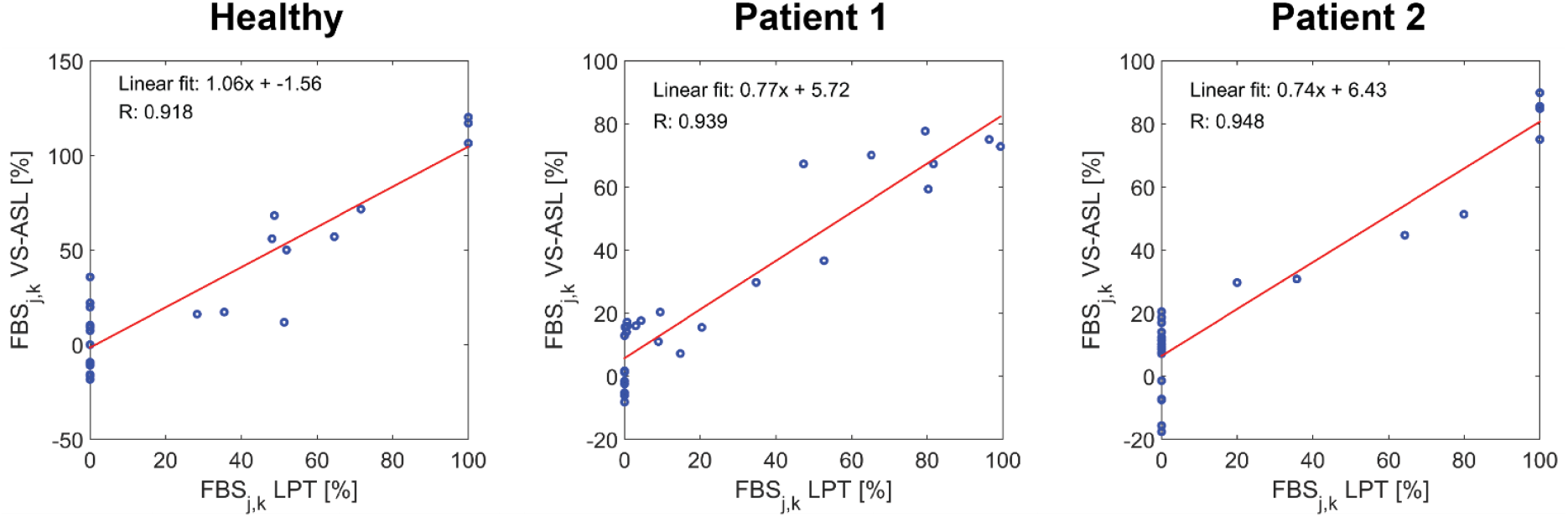
Correlation of *FBS_j,k_* between VS-ASL and LPT. For each subject, the correlation coefficient and linear fit of *FBS_j,k_* over all territories *j* and neck arteries *k* was calculated

**Fig. S3:**
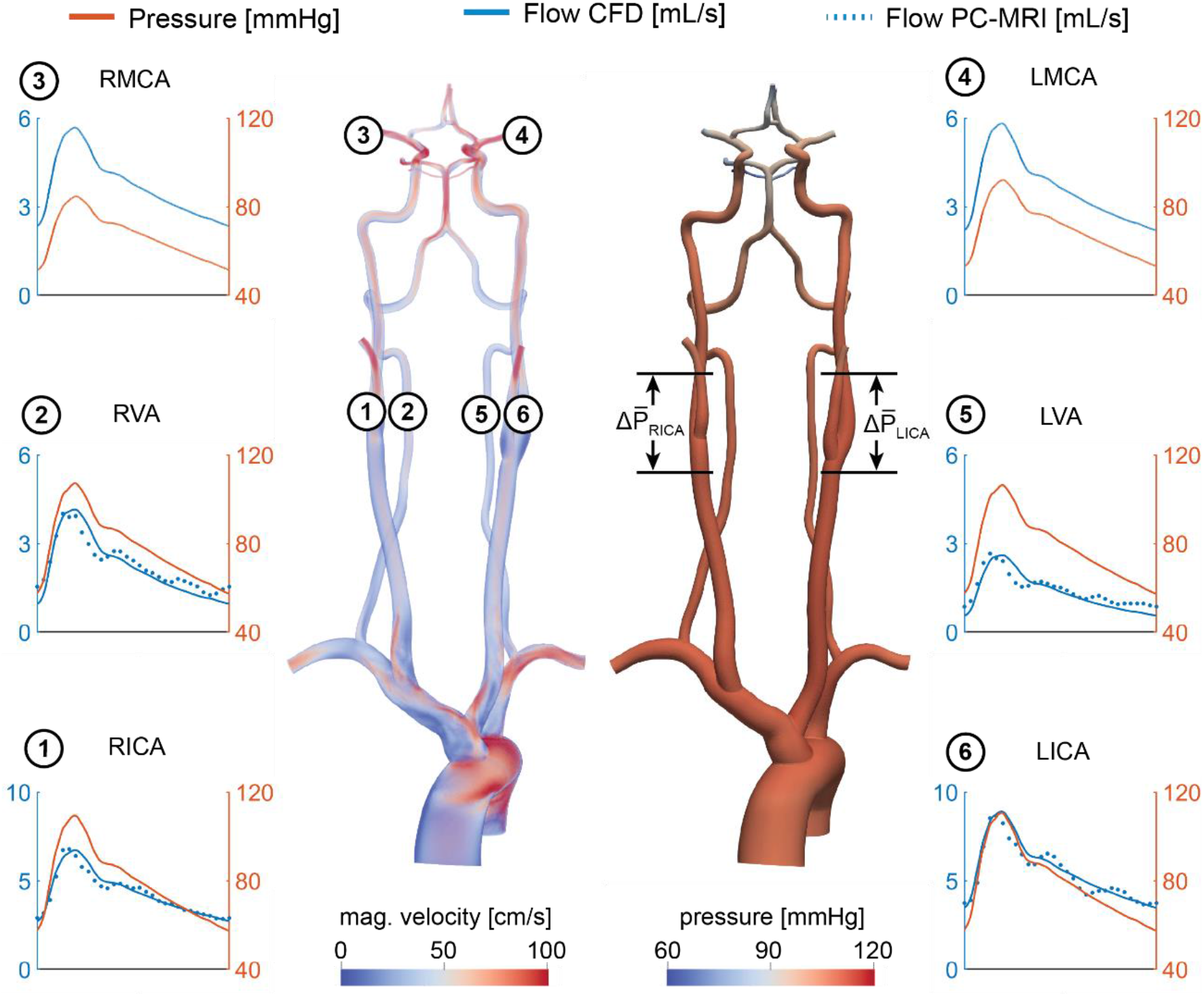
Velocity and pressure fields at peak systole are shown for the healthy control subject. Flow and pressure waveforms are evaluated in the internal carotid (ICA), vertebral (VA), and middle cerebral arteries (MCA) and compared to PC-MRI measurements in the neck arteries. The mean pressure drop 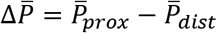 was calculated over the RICA and LICA bifurcation. The pressure and flow results for the healthy control subject are provided in the supplement material (Fig. S2). A pressure drop and fractional flow over the patent right and left carotid bifurcations were calculated analog to the patient examples for reference, yielding 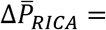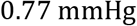 and 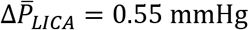 and correspondingly *FF_RICA_* = 0.99 and *FF_LiCA_* = 0.99. The flow rates in the RMCA and LMCA were similar with 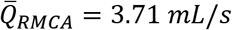 and 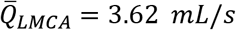 and the pressure difference between these outlets was 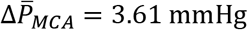.

**Table S1:** Parameters of the calibrated 3-element Windkessel models for all study subjects. Each Windkessel model consists of a proximal resistance *R_p_*, distal resistance *R_d_*, and compliance *C*. RECA/LECA = right/left external carotid artery, RACA/LACA = right/left anterior cerebral artery, RMCA/LMCA = right/left middle cerebral artery, RPCA/LPCA = right/left posterior cerebral artery, RSCA/LSCA = right/left superior cerebellar artery, RVA = right vertebral artery.

| Outlet | Healthy Control |  |  | Patient 1 |  |  | Patient 2 |  |  |
| --- | --- | --- | --- | --- | --- | --- | --- | --- | --- |
| | $R_p$<br>($10^9 \text{Pa s m}^{-3}$) | $R_d$<br>($10^9 \text{Pa s m}^{-3}$) | $C$<br>($10^{-10} \text{m}^3 \text{Pa}^{-1}$) | $R_p$<br>($10^9 \text{Pa s m}^{-3}$) | $R_d$<br>($10^9 \text{Pa s m}^{-3}$) | $C$<br>($10^{-10} \text{m}^3 \text{Pa}^{-1}$) | $R_p$<br>($10^9 \text{Pa s m}^{-3}$) | $R_d$<br>($10^9 \text{Pa s m}^{-3}$) | $C$<br>($10^{-10} \text{m}^3 \text{Pa}^{-1}$) |
| Descending Aorta | 0.01 | 0.18 | 111.52 | 0.01 | 0.23 | 63.72 | 0.01 | 0.26 | 54.52 |
| R. subclavian | 0.15 | 2.71 | 7.46 | 0.13 | 2.68 | 5.50 | 0.17 | 3.24 | 4.43 |
| L. subclavian | 0.15 | 2.70 | 7.46 | 0.13 | 2.63 | 5.50 | 0.17 | 3.24 | 4.43 |
| RECA | 0.48 | 5.14 | 3.85 | 0.49 | 5.69 | 2.44 | 0.42 | 4.90 | 2.89 |
| LECA | 0.50 | 5.30 | 3.73 | 0.85 | 10.01 | 1.42 | 0.64 | 7.46 | 1.89 |
| RACA | 3.57 | 3.32 | 2.74 | 2.38 | 2.38 | 2.09 | 4.72 | 8.01 | 1.18 |
| LACA | 3.40 | 3.15 | 2.92 | 2.11 | 2.11 | 2.42 | 5.76 | 6.23 | 1.26 |
| RMCA | 1.23 | 1.16 | 7.87 | 0.15 | 1.55 | 5.31 | 2.10 | 3.54 | 2.76 |
| LMCA | 1.35 | 1.24 | 7.70 | 1.27 | 1.27 | 5.60 | 2.61 | 2.83 | 2.86 |
| RPCA | 1.54 | 6.17 | 2.49 | 4.22 | 4.57 | 1.62 | 7.51 | 12.63 | 0.77 |
| LPCA | 1.61 | 6.42 | 2.45 | 4.37 | 4.73 | 1.55 | 8.61 | 9.33 | 0.86 |
| RSCA | 2.02 | 8.06 | 1.32 | 1.32 | 14.59 | 0.91 | 6.11 | 15.79 | 0.69 |
| LSCA | 2.51 | 10.06 | 1.32 | 1.32 | 14.03 | 0.91 | 6.14 | 15.85 | 0.69 |
| Terminal RVA | - | - | - | - | - | - | 7.09 | 17.09 | 0.61 |

## References

[1] Saba L, Yuan C, Hatsukami TS, et al. Carotid Artery Wall Imaging: Perspective and Guidelines from the ASNR Vessel Wall Imaging Study Group and Expert Consensus Recommendations of the American Society of Neuroradiology. Am J Neuroradiol 2018; 39: E9–E31.

[2] Liebeskind DS. Collateral circulation. Stroke 2003; 34: 2279–2284.

[3] Liebeskind DS, Feldmann E. Fractional Flow in Cerebrovascular Disorders. Interv Neurol 2012; 1: 87–99.

[4] Henderson RD, Eliasziw M, Fox AJ, et al. Angiographically defined collateral circulation and risk of stroke in patients with severe carotid artery stenosis. Stroke 2000; 31: 128–132.

[5] Bisschops RHC, Klijn CJM, Kappelle LJ, et al. Collateral flow and ischemic brain lesions in patients with unilateral carotid artery occlusion. Neurology 2003; 60: 1435–1441.

[6] Hendrikse J, Hartkamp M, Hillen B, et al. Collateral Ability of the Circle of Willis in Patients With. Stroke 2001; 2768–2773.

[7] Ramsay SC, Yeates MG, Lord RS, et al. Use of technetium-HMPAO to demonstrate changes in cerebral blood flow reserve following carotid endarterectomy. J Nucl Med 1991; 32: 1382–1386.

[8] Russell SM, Woo HH, Siller K, et al. Evaluating middle cerebral artery collateral blood flow reserve using acetazolamide transcranial Doppler ultrasound in patients with carotid occlusive disease. Surg Neurol 2008; 70: 466–470.

[9] Alsop DC, Detre JA, Golay X, et al. Recommended implementation of arterial spin-labeled perfusion MRI for clinical applications: A consensus of the ISMRM perfusion study group and the european consortium for ASL in dementia. Magn Reson Med 2014; 116: 102–116.

[10] Helle M, Norris DG, Rüfer S, et al. Superselective pseudocontinuous arterial spin labeling. Magn Reson Med 2010; 64: 777–786.

[11] Schollenberger J, Figueroa CA, Nielsen JF, et al. Practical considerations for territorial perfusion mapping in the cerebral circulation using super-selective pseudo-continuous arterial spin labeling. Magn Reson Med 2020; 83: 492–504.

[12] Richter X V, Helle XM, Osch XMJP Van, et al. MR Imaging of Individual Perfusion Reorganization Using Superselective Pseudocontinuous Arterial Spin-Labeling in Patients with Complex Extracranial Steno-Occlusive Disease. AJNR Am J Neuroradiol 2017; 38: 703–711.

[13] Helle M, Rüfer S, Van Osch MJP, et al. Superselective arterial spin labeling applied for flow territory mapping in various cerebrovascular diseases. J Magn Reson Imaging 2013; 38: 496–503.

[14] Jensen-Kondering U, Lindner T, van Osch MJP, et al. Superselective pseudo-continuous arterial spin labeling angiography. Eur J Radiol 2015; 84: 1758–1767.

[15] Liu J, Yan Z, Pu Y, et al. Functional assessment of cerebral artery stenosis: A pilot study based on computational fluid dynamics. J Cereb Blood Flow Metab 2017; 37: 2567–2576.

[16] Leng X, Scalzo F, Ip HL, et al. Computational fluid dynamics modeling of symptomatic intracranial atherosclerosis may predict risk of stroke recurrence. PLoS One 2014; 9: 1–8.

[17] Castro MA, Putman CM, Cebral JR. Computational fluid dynamics modeling of intracranial aneurysms: Effects of parent artery segmentation on intra-aneurysmal hemodynamics. Am J Neuroradiol 2006; 27: 1703–1709.

[18] Raschi M, Mut F, Byrne G, et al. CFD and PIV Analysis of Hemodynamics in a Growing Intracranial Aneurysm. Int j numer method biomed eng 2012; 28: 214–228.

[19] Rayz VL, Boussel L, Acevedo-Bolton G, et al. Numerical Simulations of Flow in Cerebral Aneurysms: Comparison of CFD Results and In Vivo MRI Measurements. J Biomech Eng 2008; 130: 1–9.

[20] Xiao N, Humphrey JD, Figueroa CA. Multi-scale computational model of three-dimensional hemodynamics within a deformable full-body arterial network. J Comput Phys 2013; 244: 22–40.

[21] Mukherjee D, Jani ND, Selvaganesan K, et al. Computational Assessment of the Relation Between Embolism Source and Embolus Distribution to the Circle of Willis for Improved Understanding of Stroke Etiology. J Biomech Eng 2016; 138: 81008.

[22] Bockman MD, Kansagra AP, Shadden SC, et al. Fluid Mechanics of Mixing in the Vertebrobasilar System: Comparison of Simulation and MRI. Cardiovasc Eng Technol 2012; 3: 450–461.

[23] Arthurs CJ, Khlebnikov R, Melville A, et al. CRIMSON: An Open-Source Software Framework for Cardiovascular Integrated Modelling and Simulation. Epub ahead of print January 2020. DOI: 10.1101/2020.10.14.339960.

[24] Sahni O, Müller J, Jansen KE, et al. Efficient anisotropic adaptive discretization of the cardiovascular system. Comput Methods Appl Mech Eng 2006; 195: 5634–5655.

[25] Vignon-Clementel IE, Figueroa CA, Jansen KE, et al. Outflow boundary conditions for 3D simulations of non-periodic blood flow and pressure fields in deformable arteries. Comput Methods Biomech Biomed Engin 2010; 13: 625–640.

[26] Xiao N, Alastruey J, Figueroa CA. A systematic comparison between 1-D and 3-D hemodynamics in compliant arterial models. Int j numer method biomed eng 2014; 30: 204–231.

[27] Whiting CH, Jansen KE. A stabilized finite element method for the incompressible Navier-Stokes equations using a hierarchical basis. Int J Numer Methods Fluids 2001; 35: 93–116.

[28] Kim DE, Park JH, Schellingerhout D, et al. Mapping the Supratentorial Cerebral Arterial Territories Using 1160 Large Artery Infarcts. JAMA Neurol. Epub ahead of print 2018. DOI: 10.1001/jamaneurol.2018.2808.

[29] Arthurs CJ, Lau KD, Asrress KN, et al. A mathematical model of coronary blood flow control: Simulation of patient-specific three-dimensional hemodynamics during exercise. Am J Physiol - Hear Circ Physiol 2016; 310: H1242–H1258.

[30] Di Achille P, Tellides G, Figueroa CA, et al. A haemodynamic predictor of intraluminal thrombus formation in abdominal aortic aneurysms. Proc R Soc A Math Phys Eng Sci 2014; 470: 20140163.

[31] Van Bakel TM, Arthurs CJ, Van Herwaarden JA, et al. A computational analysis of different endograft designs for Zone 0 aortic arch repair. Eur J Cardio-thoracic Surg 2018; 54: 389–396.

[32] Nauta FJH, Lau KD, Arthurs CJ, et al. Computational Fluid Dynamics and Aortic Thrombus Formation Following Thoracic Endovascular Aortic Repair. Ann Thorac Surg 2017; 103: 1914–1921.

[33] Suh GY, Les AS, Tenforde AS, et al. Quantification of particle residence time in abdominal aortic aneurysms using magnetic resonance imaging and computational fluid dynamics. Ann Biomed Eng 2011; 39: 864–883.

[34] Arzani A, Les AS, Dalman RL, et al. Effect of exercise on patient specific abdominal aortic aneurysm flow topology and mixing. Int J Numer Meth Biomed Engng 2014; 4179: 280–295.

[35] Miao Z, Liebeskind DS, Lo W, et al. Fractional Flow Assessment for the Evaluation of Intracranial Atherosclerosis: A Feasibility Study. Interv Neurol 2016; 5: 65–75.

[36] Zhang H, Fujiwara N, Kobayashi M, et al. Development of patient-specific 1D-0D simulation based on MRI and SPECT data. J Biorheol 2018; 32: 2–8.

[37] Yamada S, Kobayashi M, Watanabe Y, et al. Quantitative Measurement of Blood Flow Volume in the Major Intracranial Arteries by Using 123I-Iodoamphetamine SPECT. Clin Nucl Med 2014; 39: 868–873.

[38] Vali A, Aristova M, Vakil P, et al. Semi-automated analysis of 4D flow MRI to assess the hemodynamic impact of intracranial atherosclerotic disease. Magn Reson Med 2019; 82: 749–762.

[39] Tonino PAL, De Bruyne B, Pijls NHJ, et al. Fractional Flow Reserve versus Angiography for Guiding Percutaneous Coronary Intervention. N Engl J Med 2009; 360: 213–224.

[40] De Bruyne B, Fearon WF, Pijls NHJ, et al. Fractional Flow Reserve–Guided PCI for Stable Coronary Artery Disease. N Engl J Med 2014; 371: 1208–1217.

[41] Nouh A, Hussain M, Mehta T, et al. Embolic strokes of unknown source and cryptogenic stroke: Implications in clinical practice. Front Neurol 2016; 7: 1–16.

[42] Tariq N, Khatri R. Leptomeningeal collaterals in acute ischemic stroke. J Vasc Interv Neurol 2008; 1: 91–5.

